# Varying importance of postzygotic isolation in polyploid speciation? A survey of the triploid block strength, its causal mechanisms, and evolutionary consequences

**DOI:** 10.1101/2024.07.13.603373

**Authors:** Susnata Salony, Josselin Clo, Filip Kolář, Clément Lafon Placette

## Abstract

The triploid block, primarily caused by endosperm developmental issues, is known as a significant barrier to interploidy hybridization among flowering plants and thereby, polyploid speciation. However, its strength varies across taxa, with some instances of leakiness, questioning its universal role as a barrier. We conducted a literature survey to explore the causes of the variation in the strength of triploid block across angiosperms. We particularly assessed the impact of interploidy cross direction, types of endosperm development, endosperm persistence at seed maturity, and ploidy divergence. None of these factors had a significant impact on triploid seed viability, likely due to limited data and inconsistencies in estimation methods across the literature. In addition, triploid seed viability in experimental crosses was sometimes correlated to the occurrence of triploid hybrids in nature, sometimes not, suggesting a mixed role of the triploid block in shaping interspecies gene flow. Altogether, our study highlights the need for unified approaches in future studies on the triploid block to advance our understanding of its variation and evolutionary implications.

## 1. Introduction

Understanding the genetic basis of reproductive isolation is crucial to our understanding of the speciation process. Reproductive isolation is the result of a combination of pre- and postzygotic barriers, highlighting the importance of understanding both prezygotic and postzygotic mechanisms (Rieseberg and Willis, 2007; Lowry et al., 2008). While prezygotic barriers are typically emphasized due to their early emergence and possible prevention of costly mating (Ramsey et al., 2003; Sobel and Chen, 2014; Arnegard et al., 2014), a deeper understanding of postzygotic barriers is also crucial in speciation research (Coughlan and Matute, 2020). Here we focus on the impact of whole genome duplication (WGD) on postzygotic isolation. WGD is a dramatic mutation that leads to the production of an additional copy of the entire genome of a lineage and is recognized as a major driver of speciation (Lynch and Force, 2000; Servedio and Sætre, 2003; Rieseberg and Willis, 2007; Coughlan and Matute, 2020; Fox et al., 2020). Genome duplication occurs across Eukaryotes, though they are especially notable among plants (Cui et al., 2006). WGD is considered to bring about an instant hybridization barrier between diploids and their polyploid derivatives (Ramsey and Schemske, 1998), particularly crosses between diploid and tetraploid individuals, which is the primary focus of our review. Often these crosses result in the failure of viable seed formation, referred to as ‘triploid block’ (Marks, 1966). In the case of comparatively weaker early-acting barriers, the triploid block may emerge as a significant contributor to the genetic isolation between diploids and polyploids, and thereby polyploid speciation. However there are many studies on the genetic basis of homoploid barriers (Abbott et al., 2013; Yakimowski and Rieseberg, 2014; Nieto Feliner et al., 2017), and very little is known about the drivers and basis of heteroploid barriers in nature, despite frequent polyploid speciation (Kolář et al., 2017).

The triploid block is mostly caused by developmental problems in the hybrid endosperm (Köhler et al., 2010). The endosperm is a tissue nourishing the embryo in the seed and is the characteristic feature of most flowering plants (Baroux et al., 2002). In most diploid species, the endosperm is predominantly triploid, with two copies of the maternal genome and one copy of the paternal genome (2m:1p ratio). In interploidy hybrid seeds, this 2m:1p ratio is disturbed (Johnston et al., 1980; Haig and Westoby, 1991). Studies with artificial *Arabidopsis thaliana* polyploid series have shown that the more pronounced the deviation from the 2m:1p ratio, the stronger the defects in endosperm development and the less viable the interploidy seeds (Scott et al., 1998). With this observation followed by a large body of literature (Köhler et al., 2010; Schatlowski and Kohler, 2012; Birchler, 2014)), it is now well-accepted that the primary cause for endosperm defects in interploidy hybrid seeds is a gene dosage imbalance between maternal and paternal genomes related to the strict requirement of the 2m:1p genome ratio (Johnston et al., 1980; Haig and Westoby, 1991). This strict balance requirement suggests that paternal and maternal genomes are not equivalent, i.e., they express different genes and impact endosperm development differently. This may explain a particular feature of the triploid block, which is its parent-of-origin manifestation (Feil and Berger, 2007). Depending on whether the polyploid is the paternal or maternal parent, the phenotypic defects of the endosperm may vary as well as the rate of viability of interploidy hybrid seeds (Stoute et al. 2012; Scott et al. 1998). In crosses where the maternal parent possesses a higher ploidy level than the paternal one (referred to as ’mat-excess’ hereafter), such as in model species *A. thaliana*, the hybrids exhibit a high viability rate (Scott et al., 1998). A similar pattern has been shown in other species (Von Bothmer and Jacobsen, 1986; Sekine et al., 2013), giving the impression that mat-excess seeds generally survive better than paternal-excess ones (where the paternal parent possesses a higher ploidy level than the maternal one, referred to as ’pat-excess’ hereafter). Reports showing no parent-of-origin impact on interploidy seeds (Burton and Husband, 2000; Sonnleitner et al., 2013) question what may appear as a general trend.

Also, there is a growing realization that the triploid block is sometimes leaky, questioning how strong and universal barrier to interploidy gene flow it represents across angiosperms. Experimental crosses using a variety of natural and artificially produced diploids and tetraploids (Dinu et al., 2005; Stoute et al., 2012; Greiner and Oberprieler, 2012; Sabara et al., 2013; Sekine et al., 2013; Behrend et al., 2015; Roccaforte et al., 2015; Vallejo-Marín et al., 2016) indeed show variation in the manifestation of the triploid block across taxa, both in terms of survival rate of interploidy hybrids and its parent-of-origin asymmetry. Therefore, the triploid block may or may not be a strong barrier to interploidy hybridization, and the reason behind it is unclear. Understanding the underlying causes of this variation may help predict general patterns of interploidy gene flow in nature. However, the consequences of the triploid block on gene flow in nature are also unclear. The triploid block is one of many potential reproductive barriers preventing interploidy gene flow, and in fact, postzygotic barriers such as hybrid seed lethality are thought to play a minor role in regulating gene flow since they act late in the hybridization sequence (Coyne and Orr, 2004; Rieseberg and Willis, 2007; Lowry et al., 2008). The triploid block may therefore have little evolutionary consequences, and it is important to re-evaluate the assumption of the triploid block as an important driver of polyploid speciation.

In this review, we investigated the variation of the triploid block across angiosperms and assessed its role as a barrier to interploidy hybridization. We provide a synthesis of current knowledge to 1) evaluate how often the triploid block is a leaky barrier across angiosperms; 2) propose and test causes to explain why the strength of the triploid block varies between taxa; and 3) critically assess whether the triploid block and its variation do have an evolutionary significance in terms of realized interploidy gene flow in nature.

## 2. Variation in the strength of triploid block across angiosperms

The triploid block has been studied in several mixed-ploidy species taxa, using a variety of natural and artificially produced diploids and polyploids (Dinu et al., 2005; Stoute et al., 2012; Greiner and Oberprieler, 2012; Sabara et al., 2013; Sekine et al., 2013; Behrend et al., 2015; Roccaforte et al., 2015; Vallejo-Marín et al., 2016). In this section, we assess to what extent the strength of the triploid block shows natural variation across controlled crossing experiments. The variation in the strength of the triploid block may occur at two different levels. First, the intensity of the barrier itself might vary across taxa. Second, the cross-direction effect observed in the triploid block may vary, i.e., the viability may not always be the highest in a given cross direction (mat- or pat-excess), or even sometimes this effect may be absent, leading to equivalent triploid seed viability independent of the cross direction.

Regarding the general intensity of the triploid block, a striking range of triploid seed viability has been observed across studies. The variation spans from the virtual absence of viable seeds (0%) in crosses between diploid *Mimulus guttatus* and their natural tetraploids (Salony et al. in review) to a majority (about 80%) of triploid seeds being viable on average in *Cyclamen persicum* Mill. (Takamura and Miyajima, 1996), and anything in between. This data indicates the triploid block may be a permeable barrier to interploidy gene flow in some species, while it may represent a strong barrier in others (Fig. 3). In addition, the cross-direction effect on triploid seed viability, a commonly assumed feature of the triploid block, is inconsistent across studies. Earlier surveys of interploidy crossing data suggested that mat-excess crosses are consistently more successful than pat-excess crosses at producing viable triploid seeds (Ramsey and Schemske, 1998). Indeed, germination results often showed a higher survival upon interploidy crossing with higher maternal contribution in *Brassica oleracea*: 90% in mat-excess to complete lethality in pat-excess (Stoute et al., 2012), *Arabidopsis arenosa*: 69% in mat-excess to 7% in pat-excess (Morgan et al., 2021a), *Oryza sativa*: 50% in mat-excess to complete lethality in pat-excess (Sekine et al., 2013) or *Salpiglossis sinuata*: 8% in mat-excess to complete lethality in pat-excess (Needham and Erickson, 1992), to only cite a few (Fig. 3).

**Fig. 1.**
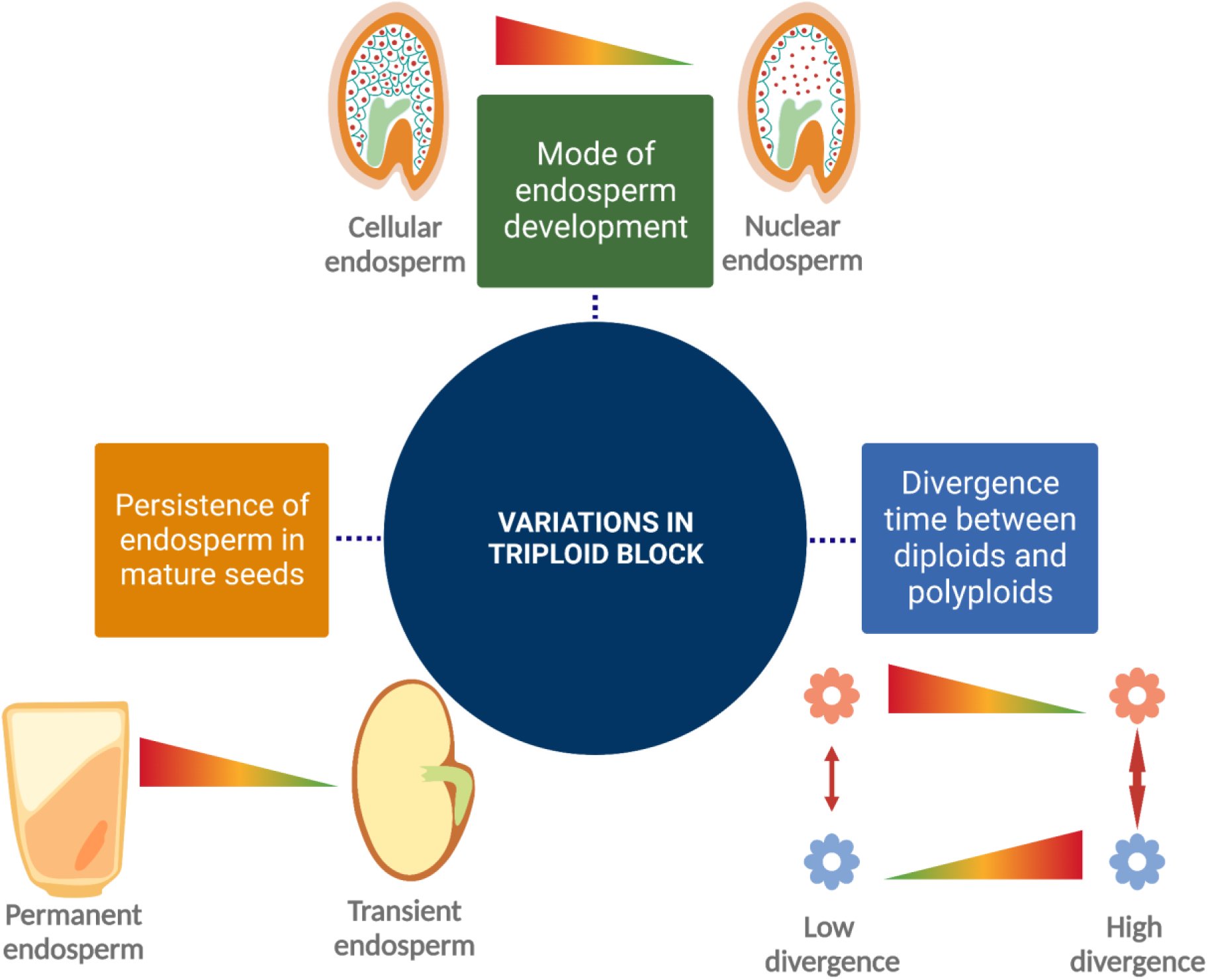
Factors influencing the variations in the strength of the triploid block, as proposed in this study. These factors include the mode of endosperm development, the persistence of endosperm in mature seeds, and the divergence time between diploids and polyploids, all of which may potentially impact the strength of the triploid block.

**Fig. 2.**
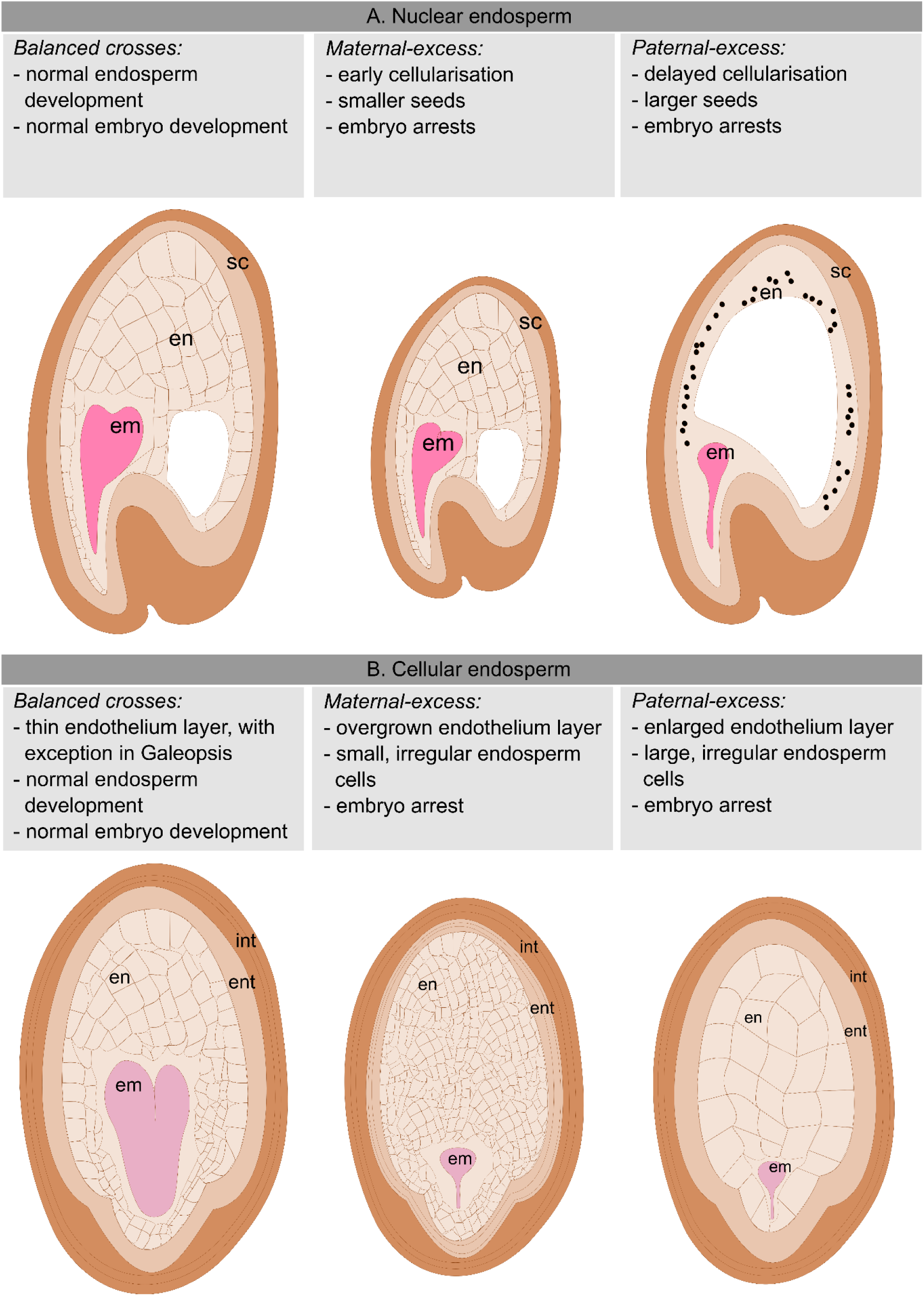
Developmental progress of endosperm, embryo and surrounding maternal tissues in the seeds obtained from control and interploidy hybridisations; a. nuclear endosperm b. cellular endosperm; em: embryo, en: endosperm, ent: endothelium, int: integumentary cells, sc: seed coat.

**Fig. 3.**
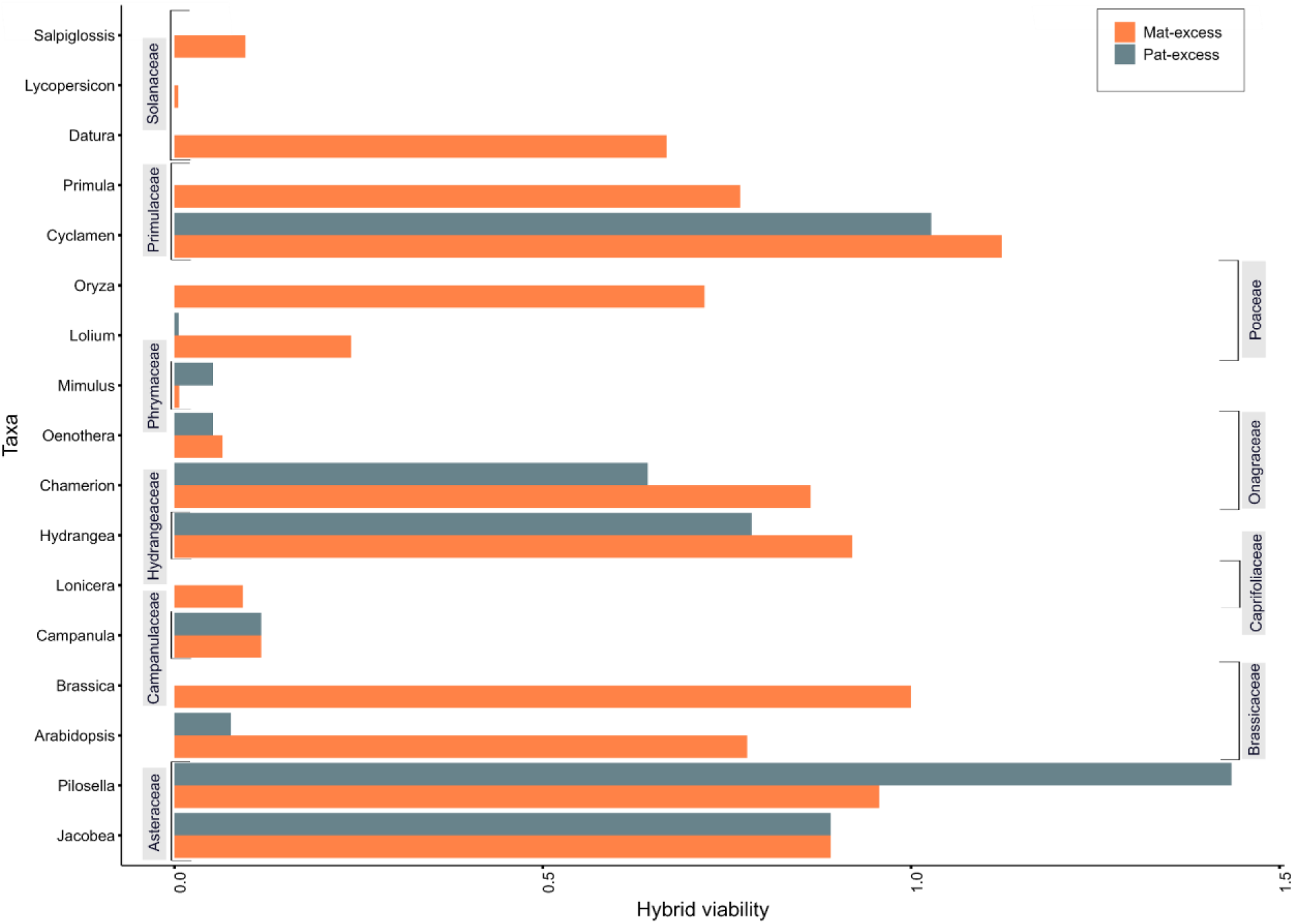
Proportion of absolute hybrid viability across species, grouped according to respective taxa studied in the present review. Mat-excess and Pat-excess hybrids are represented by red and blue respectively.

However, the commonly accepted idea that mat-excess seeds are more viable than paternal excess does not always hold. Only recently pat-excess crosses have been found to be more successful in *Mimulus guttatus* (Phrymaceae), showing an increase from 0.5% in mat-excess to 4% in pat-excess (Meeus et al., 2020). However, in Salony et al. (in review), similar experimental crosses in *M. guttatus* revealed a strong barrier to triploid formation in both directions, resulting in non-germinable seeds. Further evidence for better survival of pat-excess hybrids comes from the experimental crossings in *Pilosella echioides* (Asteraceae): 44% viability in mat-excess to 66% in pat-excess (Chrtek et al., 2017). While emasculation is not possible in Asteraceae, the high rate of viability is unlikely due to self-pollination as the species was shown to be self-incompatible in this study. While mature seeds from reciprocal crosses in *Galeopsis pubescens* (Lamiaceae) were found to be non-germinable (Hákansson, 1952), the seed viability in reciprocal crosses of other taxa, such as Solanaceae, remains unclear due to incomplete crossing designs (missing pat-excess viability) and the lack of data on seed viability (Sansome et al., 1942; Cooper and Brink, 1945; Wangenheim, 1957).

Additional evidence against the parent-of-origin effect on the viability of triploid seeds emerges from *Oenothera hookeri* (Onagraceae), where a robust triploid block is observed in both cross directions, with a viability of 5% in mat-excess and 4% in pat-excess crosses (Wangenheim, 1962). Similarly, in *Chamerion angustifolium* (Onagraceae; 69% in mat-excess/51% in pat-excess; (Burton and Husband, 2000), the triploid block is notably consistent in both cross directions, however being rather relaxed. It is interesting to note that despite similar triploid seed viability in both cross directions, the developmental defects in mat-excess and pat-excess, as studied in *Oenothera*, differ (Wangenheim, 1962). This suggests the existence of an asymmetric developmental basis for the triploid block in *Oenothera*, leading to a symmetric outcome in terms of seed viability.

It thus appears that the strength of the triploid block is quite variable across studies, ranging from acting as an absolute barrier to interploidy hybridization in some cases to being considerably permeable in other cases. This variability seems to extend to related species as well, with certain families exhibiting consistent patterns, such as Brassicaceae species showing relatively high survival rates of mat-excess seeds (Scott et al., 1998; Stoute et al., 2012; Morgan et al., 2021a), while Lamiaceae species display a robust triploid block in both cross directions (Hákansson, 1952). In essence, the strength of the triploid block may show a phylogenetic signal, with certain taxa showing a stronger triploid block than others. This leads us to propose and evaluate the potential causes for the strength of the triploid block in the subsequent section. First, the failure in endosperm development remains a consistent characteristic during interploidy hybrid seed arrest irrespective of the taxa, suggesting the developmental traits of the endosperm may influence the triploid block variations we observed above. Furthermore, differences in interploidy genetic divergence may influence the viability of hybrid seeds, as evidenced by variations in the extent of triploid block observed across primary and secondary contact zones in *A. arenosa* (Morgan et al., 2021a).

Subsequently, we aim to bring some novel insight into these hypotheses through a meta-analysis of inter-ploidy crosses conducted in the reviewed studies and uncover the underlying causes and potential consequences on gene flow and speciation.

## 3. Causes for the variation in the triploid block strength

### 3.1 Developmental differences

As introduced earlier, the triploid block is primarily manifested in the endosperm, a nourishing tissue that serves to support embryo growth (Lopes and Larkins, 1993). Failure of endosperm development ultimately causes embryo arrest and hybrid seed lethality in interploidy crosses (Scott et al., 1998; Köhler et al., 2010). Endosperm shows a wide diversity across angiosperms, ranging from its development, its persistence after seed maturation, or its ploidy (Baroux et al., 2002). The endosperm development characteristics of a given species could likely be a factor determining the extent of seed lethality upon interploidy hybridization. However, little is known about how diversity in endosperm characteristics across taxa can drive interspecific variation in the strength of the triploid block. Identifying the link between endosperm characteristics and the strength of the triploid block can serve as a basis for understanding the molecular basis of such postzygotic barriers. Therefore, we discuss here the endosperm features that could potentially influence or contribute to the variations in the strength of the triploid block. In this section, we will focus exclusively on intraspecific interploidy crosses for the sake of eliminating potential confounding factors.

#### 3.1.1 Cellular vs nuclear type of endosperm

There are mainly two types of endosperm development: nuclear and cellular. The nuclear mode of development is the most common, found in ∼ 160 angiosperm families, including 83% dicots (e.g., *Arabidopsis*, soybean, cotton) and monocots such as maize, rice, and wheat (Kordyum and Mosyakin, 2020). In nuclear endosperm formation, an initial phase of repeated free-nuclear divisions (called syncytial phase) is followed by a cellularization (cytokinesis) phase (Lopes and Larkins, 1993; Brown et al., 1999). During the syncytial phase, the endosperm central vacuole acts as the major resource sink, thereby allowing the early embryo to uptake nutrients from the surrounding endosperm mostly through supporting structures (such as the suspensor). With endosperm cellularization, there is a shrinkage of the central vacuole, thereby marking the transition to the phase when the embryo acts as the primary resource sink (additionally, the suspensor degenerates), being fed directly from the endosperm (Lafon-Placette and Köhler, 2014). Hence, the shift from the syncytial phase to the cellular phase is a crucial point (Hehenberger et al., 2012) for embryo nourishment as has been demonstrated in *Arabidopsis thaliana* mutants with defective endosperm cellularization (Kradolfer et al., 2013). Therefore, embryo arrest in response to endosperm cellularization failure may be caused by a disrupted supply of nutrients to the embryo.

Interploidy crosses in species possessing nuclear-type endosperm demonstrated deviations in the timing of cellularization in the endosperm (*Arabidopsis* - (Scott et al., 1998; Morgan et al., 2021a); *Oenothera* - (Wangenheim, 1962). In mat-excess crosses, cell wall formation is initiated earlier than the normal endosperm, leading to a shorter cell proliferation phase. This leads to smaller but plump seeds. In the reciprocal cross (pat-excess), cellularization is delayed, and as a result, cell proliferation continues longer than in normal endosperm. In addition, the central vacuole still occupies major parts of the endosperm at later stages of seed development, and hence, may remain the major nutrient sink in the seed while the embryo suffers from reduced/blocked nutrient supply (Morley-Smith et al., 2008; Hehenberger et al., 2012; Lafon-Placette and Köhler, 2014). This in turn may lead to reduced embryo growth and finally embryo arrest. In maize (*Zea mays*), (Pennington et al., 2008) reached comparable findings concerning the cellularization of endosperm and the size of hybrid seeds. It was observed that seeds resulting from mat-excess crosses were smaller as compared to those from pat-excess crosses. Moreover, the endosperm in mat-excess crosses exhibited earlier cellularization, while in pat-excess it displayed an extended period of cell proliferation. Furthermore, while the endosperm in mat-excess crosses accumulated substantial starch, in pat-excess it accumulated minimal starch due to a delayed onset of starch formation. Consequently, only a few (1.7% pat-excess, 0.83% mat-excess) seeds germinated from the reciprocal interploidy crosses in maize (Pennington et al., 2008), unlike in *Arabidopsis* where mat-excess interploidy hybrid seeds are highly viable (Scott et al., 1998; Morgan et al., 2021a). Alterations in the accumulation patterns of storage compounds were evident in rice (*Oryza sativa* - (Sekine et al., 2013) interploidy crosses, mirroring the observed pattern in maize. The endosperm from mat-excess crosses accumulated notable amounts of starch, whereas no starch granules were detected in crosses with pat-excess.

Despite a similar pattern of starch accumulation in maize and rice, they show a stark contrast in germination outcomes. Pat-excess seeds showed no germination, while mat-excess seeds exhibited high seed germination (50%) in rice. However, all resulting seedlings succumbed at an early stage. This stands in contrast to maize (Pennington et al., 2008), where the triploid block seems robust in both directions. Nevertheless, the failure of seeds involving parents with different ploidy levels can be attributed to the altered progression of endosperm cellularization-either delayed or accelerated and such aberrant endosperm development patterns may be conserved across diverse angiosperm families. It appears that the occurrence of precocious endosperm cellularization in mat-excess crosses is often less critical for seed survival compared to its absence in pat-excess crosses, as mat-excess seeds tend to exhibit better survival (i.e., seed viability) rates than pat-excess ones. However, this does not consistently follow a trend, as some taxa with nuclear endosperm deviate from this pattern.

For instance, in Rye (*Secale cereale*), the viability rates are 47.2% in pat-excess and 30% in mat-excess (Håkansson and Ellerström, 1950), and in *P. echiodies*, the rates are 66% in pat-excess crosses and 44% in mat-excess (Chrtek et al., 2017).

While the cellular mode of endosperm formation is not very common, it is found in ∼ 80 angiosperm families, mostly dicots (e.g., balsam, petunia), and only a few monocot families, such as Araceae, and Lemnaceae (Kordyum and Mosyakin, 2020). In the cellular mode of development, nuclear divisions are followed by cell-wall depositions, thus, each nuclear division (karyokinesis) is followed by a subsequent cytokinetic division, right from the beginning (Lopes and Larkins, 1993). Similar to the nuclear mode of development, the nutrient transfer in cellular endosperm takes place through adjacent maternal supporting tissues, such as the integumentary cell layers present in the seed coat (Hákansson, 1952).

During the early stages of endosperm development in *Lycopersicon pimpinellifolium* (Solanaceae), the endothelium (the inner epidermis of the integument) plays a crucial role in nutrient transfer and support for the growing endosperm and embryo (Cooper and Brink, 1945). Following fertilization, the endothelium forms a single well-defined layer of densely cytoplasmic cells surrounding the expanding endosperm. As the seed grows, the endothelium divides along the radial axis. The nutrient absorption process involves the chalazal aperture opposite the end of the conducting tissue and the endothelium. Nutrients enter the seed through the vascular bundle, moving partially through the chalazal aperture and the active cytoplasmic endothelium. Endosperm cells, especially those opposite the chalazal pocket, exhibit dense cytoplasm, suggesting an active role in nutrient absorption and hence, serving as the primary resource sink. Additionally, nutrients may diffuse into the depleted integument, being absorbed by the endosperm through the endothelium. Overall, endothelium serves as a conduit for nutrient transport, facilitating the growth and development of the seed, and any alterations in endothelium growth may be a crucial point for embryo development (Cooper and Brink, 1945; Hákansson, 1952). This has been supported by experimental crosses in *Lycopersicon pimpinellifolium* (Cooper and Brink, 1945), where the viable homoploid crosses exhibit high endosperm to endothelium ratios when compared to inviable reciprocal interploidy crosses, with smaller endosperm and thicker endothelium.

Seeds obtained from pat-excess crosses are characterized by highly vacuolated cells of endosperm and enlarged endothelial cells (Cooper and Brink, 1945). As the depletion of cells in the inner part of the integument occurs at a slower pace, there is a notable accumulation of nutrients just outside the chalazal aperture during the developmental process, thereby, blocking the nutrient supply towards the growth of the embryo. The endosperm cells in mat-excess crosses exhibit a lag in development compared to the pat-excess crosses and the seeds exhibit a smaller endosperm cavity. The endothelium cells exhibit a hyperplastic pattern, with a notable scarcity of cytoplasm in the endosperm cells adjacent to the overgrown endothelial tissue. Although accumulation of nutrients in the chalazal pocket was observed in mat-excess hybrid seeds, the embryo collapsed in the end. This suggests there could be other factors preventing the proper utilization of these reserves, leading to potential starvation of the endosperm. Although the development in reciprocal crosses proceeds quite differently and the mat-excess crosses yield significantly more mature seeds compared to that of pat-excess crosses, the seeds are non-germinable in the end (Cooper and Brink, 1945).

Similarly, no germinable seeds were obtained in *Galeopsis pubescens* (Lamiaceae) (Hákansson, 1952), although there is a pronounced difference between endosperm development in pat-excess and mat-excess crosses. In the former, the development stops very early, and degeneration sets in rapidly. In the latter, the endosperm development continues for a longer period, although at a very slow rate; but eventually degenerates. Thus, the embryo in the mat-excess cross is large compared with the small endosperm tissue, but further growth is neglectable. No clear defects of the endothelium could be observed besides a longer persistence in both pat-exc and mat-exc crosses (Hákansson, 1952). In *Datura stramonium* (Sansome et al., 1942), the rate of development of the proembryo and endosperm in the reciprocal interploidy hybrids slows down when compared to the control homoploid hybrids as the development progresses. Subsequently, the contents of the seeds disintegrate about 7 – 13 days after pollination. The disintegration may occur first in either the proembryo or in the endosperm. Endothelium cells were found to be enlarged in mat-excess crosses which eventually degenerate along with the embryo-sac contents. On the other hand, in pat-excess crosses, while the endothelium appeared as a single layer like in viable homoploid crosses, with a few large endosperm cells and some small compact endosperm cells, subsequently disintegration takes place in both proembryo and endosperm. It is interesting to note that the opening in the endothelium layer, which forms a communication between the contents of the embryo sac and the outer maternal tissue at the chalazal end, was still evident in the inviable interploidy crosses. However, the seed contents degenerate, thereby, suggesting a poor utilization of the nutrient reserves, leading to potential embryo arrest. Only 0.5 % of seeds were germinable in mat-excess crosses, while in pat-excess crosses, germinable seeds were rarely obtained.

To conclude, for the cellular type of endosperm, changes in the somatic tissue of the seed, especially the maternal endothelium tissues surrounding the embryo sac, and the altered development of endosperm, may cause an interrupted nutrient supply to both the endosperm and the embryo. The mechanism through which such altered development leads to the observed disintegrations, or the involvement of other factors, remains unknown at present, despite the common incidence of the cellular mode of development. In our study, we did not observe a clear trend of mat-/pat-excess asymmetry concerning triploid seed viability, unlike in nuclear endosperm. Both mat- and pat-excess crosses exhibited low survival rates, despite showing significantly different developmental courses.

#### 3.1.2 Transient vs permanent endosperm

In many species, the endosperm is consumed before seed maturation (transient endosperm) (Vijayaraghavan and Prabhakar, 1984; Becraft et al., 2001; Simpson, 2010). However, it is persistent (permanent endosperm) in others, such as cereal grains where the endosperm stores the seed reserves and represents a major source of food and industrial value for human nutrition (Vijayaraghavan and Prabhakar, 1984; Becraft et al., 2001; Simpson, 2010; Awika, 2011). In permanent endosperm, the embryo continues to use nutrients from the endosperm during seed germination, while in the transient endosperm, the endosperm ceases to exist and instead cotyledons represent the major reservoir for nutrients required during seed development and germination (Yan et al., 2014). Transient endosperms are often a characteristic feature of dicot crops such as soybeans and peas, and model plants such as *Arabidopsis* (Li, 2017).

In rice and maize (Pennington et al., 2008; Sekine et al., 2013), virtually no interploidy seedling could be retrieved. This was due to non-germinable seeds to a large extent, but interestingly, in rice, while mat-excess seeds could germinate at relatively high rates (50%), seedlings died at an early stage. This may reflect the role of the endosperm in supporting the embryo after germination, and its impaired function in interploidy seeds.

Similarly, reciprocal hybrids in *Galeopsis* and *Oenothera* are barely viable (Hákansson, 1952; Wangenheim, 1962). However, in taxa such as *Cyclamen* and *Pilosella* (Takamura and Miyajima, 1996; Chrtek et al., 2017), reciprocal interploidy crosses exhibit relatively higher hybrid viability, despite these species sharing the common characteristics of having a permanent endosperm. On the other hand, in taxa with a transient endosperm, the variation also ranges from mostly viable seeds in crosses with higher maternal dosage, as observed in *Arabidopsis* and *Brassica* (Scott et al., 1998; Stoute et al., 2012) to the near absence of viable seeds in reciprocal crosses in *Mimulus* (Meeus et al., 2020); Salony et al. in review).

To conclude, cereal crop species appear more sensitive to endosperm defects during interploidy hybridization. This sensitivity cannot be solely attributed to the persistence of the endosperm, but may be influenced by the characteristics of the family and other factors that reinforce the triploid block. We also hypothesized that taxa such as Brassicaceae or Fabaceae, which have transient endosperms, might better survive endosperm defects as this tissue appears to play a role only during a restricted developmental window, which we tested further in this study (Chapter 3.3).

### 3.2 Genetic divergence between diploids and their polyploid relatives

Triploid block is a postzygotic barrier, primarily caused by a dosage imbalance between the maternal and paternal genomes in the endosperm. However, endosperm-based hybrid seed lethality also occurs between species of same ploidy (Städler et al., 2021) as a result of negative epistasis between mutations that arose independently in each species, following the Bateson-Dobzhanksy-Meyer incompatibility model (Orr, 1996). In this model, the probability of incompatible mutations increase exponentially with divergence time between lineages. It is therefore likely that the divergence time between diploid and polyploid lineages will affect the survival of interploidy hybrid seeds. In other words, we can expect that the more genetic divergence between diploids and polyploids, the more time to accumulate incompatible mutations and thus the stronger the manifestation of the triploid block.

Alternatively, after WGD, polyploid genomes gradually decrease, through genome downsizing and the loss of duplicated genes and non coding regions (Buggs et al., 2009; Wendel et al., 2018). These changes lead to a gradual reduction in the genome size in polyploid individuals, which could weaken the triploid block by reducing gene dosage imbalances and skewing gene expression patterns towards the diploid-like state. This is likely to be more prominent in “older” polyploid lineages than younger ones.

Altogether, genetic divergence between diploids and polyploids is likely to have conflicting effects on the strength of the triploid block depending on the specific polyploid system and the age of the polyploid lineage. However, these hypotheses remain to be tested empirically, which is the focus of the subsequent section.

### 3.3 Testing of the hypotheses with a meta-analysis

To go beyond mere literature review and speculation, we tested formally with a meta-analysis whether the type of endosperm (cellular/nuclear; transient/permanent) or the divergence between diploids and polyploids could be associated with the strength of the triploid block or with the parent-of-origin pattern of hybrid seed viability. While we could directly assign a type of endosperm to each species based on the literature, we had to rely on a proxy to assess the divergence between diploids and polyploids. We compared synthetic and natural polyploids, assuming that synthetic polyploids are (nearly) isogenic while natural polyploids already evolved independently from diploids, therefore synthetics are genetically closer to diploids than natural polyploids.

#### 3.3.1 Compilation of the dataset

We reviewed 29 published studies of mixed-ploidy species in the literature that comprised diploids and natural auto-tetraploids and involved interploidy crossing data to address the variation in the strength of the triploid block and identify a common pattern, if any. We performed the literature survey using Google Scholar, using the keywords [“triploid block” or “triploid hybrids” or “interploidy hybrids” or “interploidy crosses” or “hybrid seed failure”] and [tetraploids] and [plants] from the 1900s to the present. A given study was included in our dataset only if (1) the focus was comparing diploid plants with their tetraploid counterparts from the same species (intraspecific crosses) (2) either or both of the diploid or tetraploid control estimates for hybrid seed viability was measured (3) either or both of the pat- or mat-excess estimates for hybrid seed viability was measured (4) strength of triploid block was estimated by phenotyping of mature seeds (and not ovules) (5) germination or seed abortion rates were used as a proxy for seed viability. The proportion of seeds that appeared viable was used as a proxy when germination rate data was unavailable, provided there was a proper description of seed morphology. In most papers, it was possible to extract numerical values from tables, but with a single exception, where we extracted the data directly from the figures using ImageJ (Schneider et al. 2012). The selected studies generally reported hybrid seed viability as either the germination proportion or morphologically described and categorized the viable and inviable seeds. In the later cases, we computed the viability proxy by taking a ratio of viable seeds to the sum of viable and inviable seeds. Rarely, some studies reported hybrid seed viability data using multiple replicates of control and reciprocal interploidy hybridizations. In such cases, we considered the average value for each hybridization treatment.

In total, we reviewed 19 angiosperm species representing 19 genera from 11 families (see Supplementary Table 1). Three species belong to Poaceae and Solanaceae; four families are represented by two species, and five families are represented by a single species (Χ² = 190, df = 180, p = 0.2903) (see Supplementary Table 1). Of the 19 species examined, 11 exhibited a nuclear type of endosperm and 8 exhibited a cellular type. In total, we have 31 estimates for hybrid viability from diploid (N = 15) and tetraploid (N = 16) control crosses (the remaining values were not available, indicated as NA), while 16 and 19 estimates respectively for seed viability of pat- and mat-excess hybridizations. It’s crucial to emphasize that we obtained estimates inconsistently distributed across species when considering all the factors examined to account for variations in the triploid block strength. While having all four estimates (seed viability in 2x-control, 4x-control, pat-excess, and mat-excess crosses) from a single species would enhance the accuracy of gauging the triploid block’s strength for that species, unfortunately, this is not the case (see Supplementary Table 1).

#### 3.3.2 Statistical analysis

We chose to perform a Bayesian meta-analysis by using the MCMCglmm package (Hadfield, 2010). We wrote the following model:

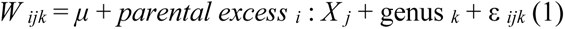

where W ijk is the fitness of triploids, μ is the mean value, and parental excess i is the effect of the parental excess i (pat-excess or mat-excess), in interaction with another fixed factor X j being potentially

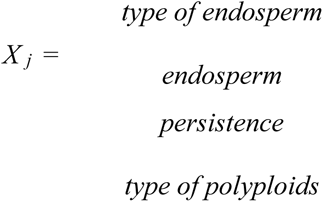

In these factors, the type of endosperm can be cellular or nuclear, the persistence of endosperm in mature seeds can be transient or permanent, and the polyploid can be either natural or synthetic. We only included a single random effect: genus k is the effect of the kth genus, and εijk is the residual error. We chose to control the variability among genera rather than among species (which is generally the case in meta-analyses). We assumed that the residual error followed a Gaussian distribution. We performed these models with two different datasets, one using the relative performance of triploid individuals compared to parental lines, and the other using the absolute viability of triploids. The relative performance of mat-excess or pat-excess triploids was determined by taking the ratio of absolute values of the respective interploidy hybrid to the average value of the homoploid hybrids.

For all analyses, we used the weakly informative, default priors proposed in MCMCglmm (Hadfield, 2010). For fixed effects, the prior is a normal distribution with the mean being equal to zero and a variance of 10^10^. For random effects, inverse-Wishart priors are implemented, with the degree of belief parameter being equal to zero and the expected variance being equal to 1. For all models, we used a burn period of 1.000.000 iterations, with a thinning interval of 50, and the MCMC chains were run for 6.000.000 iterations in total.

The parameter models and associated 95% credible intervals were thus inferred from the sampling of the posterior distribution 100.000 times. We did a visual examination of the convergence, posterior traces, and autocorrelation values of our models, as suggested in (Hadfield, 2010).

#### 3.3.3. Results

As the analyses based on the relative and absolute hybrid seed viability gave similar results, we chose to only present the results for the relative performance. None of the factors we tested were significant (Fig. S1). We found a consistent trend of mat-excess hybrids being more viable than the pat-excess ones, but no significant differences have been found (Fig. 4).

**Fig. 4.**
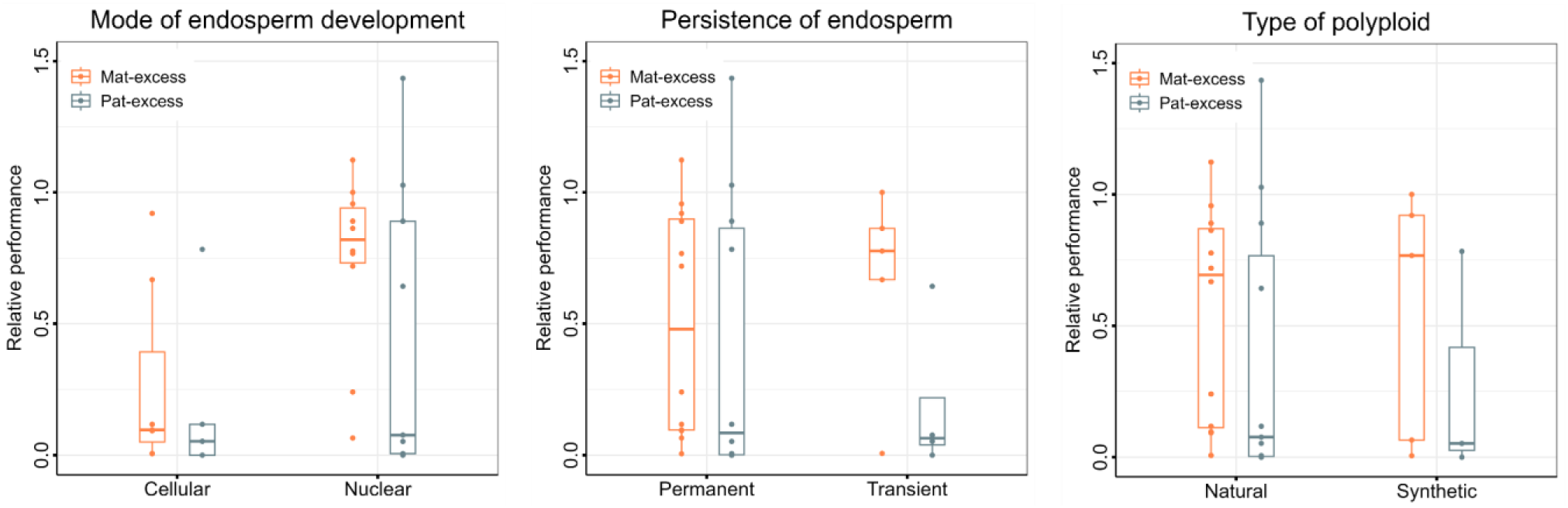
Asymmetry in the relative performance of triploid hybrids (seed germination) with respect to control homoploid hybrids, tested against different factors that were hypothesized to potentially affect the triploid embryo viability: type of endosperm development *(left)*, persistence of endosperm in mature seeds *(middle)*, type of polyploids used in interploidy crosses (natural or synthetic) *(right)*. Mat-excess and Pat-excess hybrids are represented by red and blue respectively.

We found no differences between species having cellular or nuclear endosperm (Fig. 4, *left*), no differences between species having transient or permanent endosperm (Fig. 4, *middle*), and finally no differences between triploids involving synthetic or natural polyploids (Fig. 4, *right*).

#### 3.3.4. Discussion

In this section, we investigated the impact of various aspects of endosperm development and the degree of genetic divergence between ploidies on the variation of the triploid block strength. However, none of the factors we tested demonstrated a significant effect on the strength of the triploid block. Notably, Brassicaceae plants exhibited a pattern of strong mat-excess and weak pat-excess triploid block, but this trend was not statistically significant.

Nevertheless, it is crucial to acknowledge that the limited number of studies included in our analysis makes it difficult to draw definitive conclusions about the factors influencing triploid block strength. The mechanistic basis of triploid block is primarily studied in species possessing nuclear endosperm including cereal crops and the endosperm-embryo interaction in heteroploid crosses is limited to model species, such as *A. thaliana*. Consequently, the intricate interplay of the triploid block in cellular endosperm remains poorly understood, with a lack of available information, which makes it hard to link triploid block strength and development causes for this type of endosperm. Hence, there is an urgent need to explore the strength of the triploid block in non-model species to gain deeper insights into the general causes of large variations in triploid block strength across angiosperm diversity.

Furthermore, there is a significant lack of consistency and coherence among studies concerning the estimation of triploid hybrid seed viability. Relying solely on heteroploid crosses without including homoploid crosses as control can result in an incomplete and inaccurate assessment of the triploid block strength. However, many studies did not include control homoploid crosses in their investigation, limiting the power of their inference of the triploid block strength for multi-species comparisons. In addition, the lack of a unified approach in scoring triploid viability, particularly scattered over-assessments of ovules and mature seeds or utilizing seed sets or even fruit sets as a proxy, further hampers an accurate estimation. Therefore, there is a pressing need for a comprehensive and standardized method to estimate the strength of the triploid block, one of them being simply seed germination with sufficient sample size (>100 seeds per cross), which was, unfortunately, lacking in many studies we found. Following such a standardized approach and conducting rigorous comparative analyses can provide a better understanding of the cause for the variation of triploid block strength across angiosperms.

## 4. Evolutionary consequences of the triploid block

The range of variation in the strength and parent-of-origin pattern of the triploid block may bear consequences for interploidy gene flow in nature. For example, taxa with leakier triploid block may show more pronounced gene flow compared to taxa with a strong triploid block. The real manifestation of the triploid block, however, depends on the strength of the other barriers acting prior (prezygotic) or after (later postzygotic) triploid block. While prezygotic barriers may have a key role in restricting gene flow (Husband and Sabara, 2004), postzygotic barriers such as seed inviability are presumed to have a minor role in building reproductive isolation since they act late in the hybridization sequence (Coyne and Orr, 2004). In this section, we evaluate how well we can predict genetic isolation between ploidies by comparing data on the triploid block from experimental crosses with the measure of interploidy gene flow in nature.

One way to assess the strength of interploidy isolation is to measure the presence of triploid hybrids in natural populations. The occurrence of triploid hybrids largely varies depending on the species, ranging from a large number of triploids in *Pilosella echiodies* (up to 73%; (Trávníček et al., 2011) or *P. rhodopea* (up to 58%; (Šingliarová et al., 2011), or *Galax urceolata* (up to 23%; (Burton and Husband, 1999), to the virtual absence of such hybrids in other systems such as *Arabidopsis arenosa* (0.14%; (Morgan et al., 2020) or *Mimulus guttatus* (0, Salony et al. in review). The variation in the occurrence of triploid hybrids in nature could be underlain by variations in the strength of the triploid block between taxa. However, it could also be the result of other factors, such as clonal propagation promoting the propagation of the odd-ploidy cytotypes (Trávníček et al., 2011; Chrtek et al., 2017), or a large set of prezygotic barriers preventing any interploidy hybridization in the first place. It is therefore important to evaluate whether the triploid block or other factor(s) are most likely to explain the range of triploid hybrid occurrence observed in different systems.

Although numerous natural cytotype surveys have been done in mixed-ploidy plants, and numerous studies estimated the triploid block strength with experimental crossings, very few studies have actually combined both in an attempt to relate the triploid block strength with the presence of triploid hybrids in nature. In *Chamerion angustifolium*, a species with relaxed triploid block (Burton and Husband, 2000), a significant proportion of triploids (9%) were found in natural populations (Husband and Schemske, 1998). Also, we see a strikingly high triploid frequency in *Pilosella echiodies* (or *Hieracium echiodies*) in the field (around 73%; (Trávníček et al., 2011), which corroborates the higher triploid viability (averaging 77 % in both directions) found in a crossing experiment performed by (Chrtek et al., 2017). On the contrary, *Plantago media* exhibits robust triploid block, as evidenced by the rare triploid formation in controlled crossing experiments (0.8% triploid hybrids; (Van Dijkt and Van Delden, 1990). Consistently, triploids were also rare in the mixed ploidy populations (0.44%, (Dijk et al., 1992). Similarly, the triploid block was found to be strong in *Oenothera hookeri* (approx. 5% survival; (Wangenheim, 1962), and no natural triploids were observed in this species. From the above studies, one may propose that there is at least a correlation between the triploid block and the occurrence of triploid hybrids in nature in the extreme cases: when the triploid block is strong, logically, (nearly) no triploids are reported from the field. On the other hand, frequent triploids are found in some species with known relaxed triploid block.

There are, however, cases where experimental data do not correspond with field observations. For example, in *Arabidopsis arenosa*, mat-excess crosses result in largely viable (∼75%) triploid progeny (Morgan et al., 2021a), while triploid plants are virtually absent from natural mixed-ploidy populations (Morgan et al., 2020), likely due to strong prezygotic post-pollination barriers (Morgan et al., 2021b).

Overall, while reciprocal crosses in interploidy hybridization experiments provide a valuable understanding of reproductive barriers, they may not necessarily translate to similar patterns in nature due to the complex interplay of reproductive isolation pathways in natural populations. However, studies comparing the manifestation of the triploid block after experimental crosses with interploidy gene flow in nature are too rare to firmly conclude on this point. Future interdisciplinary investigations of the mechanism (crossing experiments) and realized impact in the field (population genetics and cytogeography) of triploid block may provide valuable insights into the implications for reproductive isolation and evolutionary processes.

## 5. Conclusions & perspectives

Understanding the strength of the triploid block is of significant importance as it has implications for speciation and the dynamics of polyploid populations. Triploid block can act as a reproductive barrier between different ploidy levels, contributing to speciation. If the triploid block is strong and effectively prevents the formation of triploid hybrids, it can promote the formation of distinct diploid and polyploid lineages, thereby leading to the establishment of new species over evolutionary time. In contrast, an incomplete triploid block may allow the formation of triploid hybrids, leading to increased levels of hybridization and gene flow between different ploidy levels. This can result in higher levels of genetic diversity within polyploid populations and potentially contribute to the generation of evolutionary novelties with adaptive advantages and may promote neo polyploid establishment.

Our present study shows that the triploid block is not always a strong barrier to interploidy hybridization. Instead, the strength of the triploid block largely varies depending on the species, suggesting different outcomes for interploidy hybridization in nature potentially due to mere phylogenetic signal. We tested the impact of types of endosperm development and genetic divergence between diploids and polyploids, but could not identify any clear cause for this variation. We also could not provide statistical support for the commonly accepted idea that mat-excess seeds survive to a better extent than pat-excess ones. While this could suggest that previous hypotheses related to the triploid block mechanisms are wrong, a likely explanation is the lack of suitable and comparable data available in the literature. Despite a large number of studies with interploidy experimental crosses, too few have actually robust sample sizes, parental controls, or clearly defined criteria to assess basic proxies of triploid block such as seed viability. This calls for future works on the triploid block in a unified and rigorous setup, which should include: 1) a clear definition of what “viable seed” is assessed, and seed germination should be the ultimate measure for seed viability; 2) control crosses for both the diploid and the polyploid parent, as many studies we encountered either had no control crosses or only one of the two parents; 3) a clear definition of “seed set” as the total number of seeds (including inviable ones); when defined, “seed set” sometimes referred either to the total number of seeds or to the number of viable-looking seeds. The first definition measures prezygotic isolation while the second confounds both pre and postzygotic isolation. Finally, the impact of the triploid block on gene flow in nature remains questionable, even though, again, too few studies are available to draw firm conclusions on this matter.

## Funding

This work was supported by the Czech Science Foundation (project 20-22783S to FK) and Charles University Research Centre program no. PRIMUS/19/SCI/02 to CLP. Additional support was provided by a student grant of the Charles University Grant Agency (GAUK project no. 321222 to S.S.).

## Author contributions

C.L.P., S.S. and F.K. conceived the study. S.S. conducted the literature search. S.S. and J.C. carried out data analysis. S.S., J.C., and C.L.P. wrote the manuscript. S.S., J.C., F.K and C.L.P. reviewed and edited subsequent versions of the manuscript. All authors gave final approval for publication.

## Acknowledgements

The authors acknowledge the valuable contributions of Anna Nowicka and Aleš Pečinka in providing triploid block data regarding Barley and for their feedback on improving the manuscript.

## Declaration of interests

The authors declare no competing interests.

## Supplemental information

**Fig. S1.**
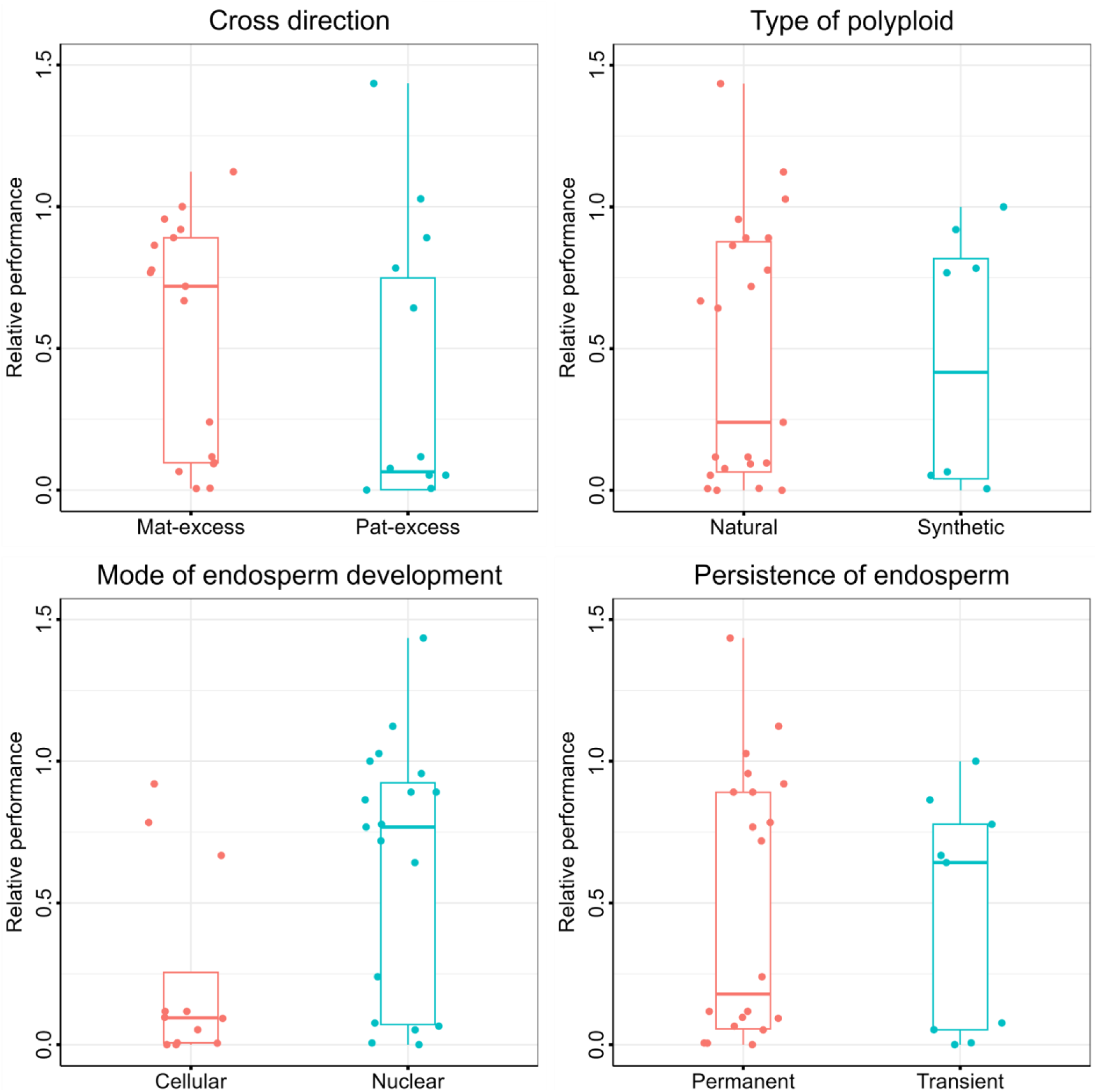
Relative performance of triploid hybrids wrt different factors that could potentially affect the triploid viability: asymmetry in parental genome contribution (during reciprocal interploidy crosses) *(top left)*, type of polyploids *(top right)*, type of endosperm development *(bottom left)*, persistence of endosperm in mature seeds *(bottom left)*.

## Notes

### Competing Interest Statement

The authors have declared no competing interest.

